# Insights into Aspergillus fumigatus morphogenesis and pathogenesis through the putative lipid transporter ArvA

**DOI:** 10.64898/2025.12.04.692327

**Authors:** Cecilia Gutierrez-Perez, Jane T. Jones, Charles T.S. Puerner, Sandeep Vellanki, Nicole E. Kordana, Matthew R. James, Elisa M. Vesely, Angus Johnson, Robert A. Cramer

**Author notes:** Corresponding Author: Robert A. Cramer, Ph.D. Authors contributed equally to this work (Cecilia Gutierrez-Perez, Jane T. Jones).

## Abstract

*Aspergillus fumigatus* poses a significant threat to human well-being, in part due to the increasing emergence of strains resistant to frontline antifungal therapy. In this study, we observe that the gene, *arvA*, is required for *A. fumigatus* morphogenesis, antifungal drug susceptibility, and cell wall homeostasis. Intriguingly, our study reveals novel morphological and growth aberrations in the absence of *arvA*. Loss of *arvA* results in hyper-swollen conidia that give rise to stunted, polarity-deficient hyphae in numerous environmental conditions, indicating a pivotal role for *arvA* in *A. fumigatus* morphogenesis. Surprisingly, despite these severe *in vitro* morphological and cell wall defects, *arvA* was not required for morbidity and mortality in immunologically distinct murine models of invasive pulmonary aspergillosis (IPA). However, growth in natural calf lung surfactant was able to normalize *ΔarvA* growth with the wild-type strain suggesting lung surfactant may partially complement the severe *in vitro* morphological defects of *arvA* loss *in vivo*. Taken together our observations reveal *arvA* as a mediator of *A. fumigatus* antifungal drug susceptibility and highlight the complex and ill-defined pulmonary nutrient environment’s role in mediating *A. fumigatus* pathogenesis and disease progression.

**IMPORTANCE:** *Aspergillus fumigatus* is a challenging fungal pathogen in the clinic in part due to increasing azole drug resistance. In this study, we observe that loss of the *A. fumigatus* gene *arvA* results in increased azole susceptibility and significant *in vitro* morphological changes highlighted by hyper-swollen conidia that yield stunted and polarity deficient hyphae.

Importantly, despite these severe *in vitro* morphological and growth abnormalities, *ΔarvA* surprisingly retains full pathogenicity and virulence in two immunologically distinct murine models of invasive pulmonary aspergillosis. These results challenge our understanding of the in-host environment and how it mediates fungal morphogenesis and pathogenesis. These results, consequently, not only enhances our understanding of the role of *arvA* in *A. fumigatus* morphogenesis and drug susceptibility, but further emphasizes the importance of *in vivo* animal models in fully evaluating potential antifungal drug targets.

## INTRODUCTION

*Aspergillus fumigatus* is a filamentous fungus that can be a pathogen in many hosts with compromised or altered immune system function. In 2024, *A. fumigatus* was estimated to cause upwards of 2 million cases of invasive aspergillosis per year worldwide with a crude mortality of ∼85% (1). Invasive aspergillosis clinical presentation is highlighted by extensive hyphal growth that often results in the formation of biofilms in tissue. Hyphae and associated biofilms present a unique challenge to not only the immune system but also contemporary antifungal therapy treatments (2). Consequently, studies of hyphal morphogenesis and biofilm development are important to identify mechanisms amenable to *in vivo* relevant antifungal therapeutic development.

The ER-resident protein Arv1 functions in ceramide, sterol and GPI-anchor precursor transport, sphingolipid metabolism, ER integrity, fatty acid resistance, hyphal formation, virulence, autophagy, and cell cycle progression, all processes critical for biofilm development (3–11). The *ARV1* gene is conserved among eukaryotic species, with its physiological function as a putative lipid transporter conserved between pathogenic and non-pathogenic fungi (12). Arv1 has been extensively characterized in *Saccharomyces cerevisiae*, *Candida albicans,* and in the context of human disease (3–6, 9–20), but its role in *A. fumigatus* hyphal morphogenesis and pathogenicity is not defined.

In *C. albicans,* Arv1p is necessary for proper sterol distribution, hyphal growth, virulence, and azole sensitivity (5, 6). Consistent with *S. cerevisiae* synthetic lethal analyses, loss of *C. albicans ARV1* results in azole hyper-susceptibility. Increased azole susceptibility in the absence of *ARV1* is likely due to Arv1p forming a heterodimer with Erg11p, the target for azole drugs, to stabilize Erg11p and increase its half-life (6, 21). Arv1 mutants that were unable to interact with Erg11p were avirulent and hyper-susceptible to azoles, suggesting that this interaction is necessary for causing disease *in vivo* and azole susceptibility (6). Loss of Arv1p also results in Erg11p mis-localization (6). In *S. cerevisiae*, loss of *ARV1* results in defective ceramide transport from the ER to the Golgi which infringes on sphingolipid metabolism (3, 4).

Furthermore, synthesis of GPI-anchors is compromised in *Δarv1* mutants (3). Arv1p interacts with GPI biosynthetic machinery and has been postulated to be a GPI flippase (15, 22). *Δarv1* has similar phenotypes to GPI-anchor synthesis mutants such as increased accumulation of chitin, hyper-susceptibility to the chitin inhibitor calcofluor white, and accumulation of GPI-anchor precursors (3, 23). In *S. cerevisiae, Δarv1* cells over-accumulate sterols in the ER while lacking sterols in the plasma membrane (4, 11). As a result, ER integrity and morphology is compromised in *Δarv1* leading to increased UPR activation (10). This result is consistent with Arv1p’s hypothesized role as a retrograde sterol transporter.

Arv1’s roles in lipid homeostasis and GPI anchor synthesis are conserved in human cells (11, 13, 14, 17). Outside the roles reported in fungi, Arv1p contributes to cell cycle progression in human cells (8). Arv1p can recruit multiple proteins necessary for cell cycle progression to the cleavage furrow, including myosin (8). Lack of Arv1p resulted in delayed telophase progression and an increase in multinucleate cells (8). These phenotypes are ill-defined in fungi but are suggested to be independent of Arv1p’s roles in lipid homeostasis (8).

In this work, we identified the *Aspergillus fumigatus* Arv1 homolog, herein named *arvA*. We hypothesized that loss of *arvA* would impact hyphal morphogenesis, biofilm formation, antifungal drug susceptibility, and virulence in a mouse model of Invasive Aspergillosis. Loss of *arvA* in *A. fumigatus* generated phenotypes similar to those observed in other fungal species upon Arv1p loss, such as azole susceptibility, delay in hyphal formation, changes in cell wall composition, and an inability to grow at high temperatures. However, loss of *arvA* in *A. fumigatus* revealed distinct and previously unreported morphological changes in agar-based and submerged biofilms *in vitro*. Strains lacking *arvA* are unable to form hyphae that grow radially in agar-based colony biofilms and instead grow as compact micro-colonies. In a submerged biofilm model, Δ*arvA* conidia are hyper-swollen and multinucleate, with a loss of polarity and conidia reaching a size 10-12 times larger than wild-type and reconstituted strains. Surprisingly, in spite of *ΔarvA’s* striking stunted morphology *in vitro*, *ΔarvA* is able to grow *in vivo* in the mouse lung and has full virulence as measured by murine mortality in two immunologically distinct invasive pulmonary aspergillosis murine models. We observe that *in vitro* growth in natural calf lung surfactant is able to partially complement the morphology and growth defects associated with loss of *arvA*, which suggests that the carbon/nutritional environment in the lung normalizes growth and virulence in the absence of *arvA*. These results highlight the gap between *in vitro* laboratory conditions and the physiological conditions *in host*.

## RESULTS

### Loss of function of arvA (AFUB_027230) results in striking morphological changes compared to parental strain CEA10

We became interested in the *A. fumigatus* Arv1p homolog, AFUB_027230 (here in called *arvA* consistent with *A. fumigatus* gene nomenclature convention), due to its synthetic lethal interaction with Erg11p in *S. cerevisiae* (*24*). Loss of function of *S. cerevisiae* Arv1 increases susceptibility to alkaline pH, azoles, statins, allylamines, and calcofluor white (6, 14, 25–27).

ArvA is a likely ortholog to human ARV1 (BLASTp 23% query cover with 33.02% identity) and *S. cerevisiae* Arv1p (BLASTp 16% query cover with 50.7% identity) as indicated by reciprocal BLAST queries. ArvA also has the predicted canonical ARV1 domain. Subsequently we generated an ArvA null mutant strain *(ΔarvA*) and reconstituted strain (*ΔarvA + arvA)* using laboratory strain CEA10 as the parental strain and a CRISPR/Cas9 mediated approach (primers indicated in **Table S1**).

Following confirmation of gene replacement and reconstitution, isogenic strain colony biofilm growth was observed on glucose minimal medium (GMM) Petri plates incubated for 72 hours at 37°C, 5% CO_2_ (**Fig. 1A**). After 72h, *ΔarvA* exhibited a significant decrease in hyphal growth as evidenced by a dramatic decrease in colony diameter compared to strains CEA10 and *ΔarvA+arvA* (**Fig. 1A**). The parental (CEA10) and reconstituted strain (*ΔarvA+arvA)* grow to about 50 mm colony diameter after 72 hours, while *ΔarvA* only reaches 4.8mm (**Fig. 1A**). This result suggests ArvA is necessary for normal colony biofilm development in commonly used laboratory *A. fumigatus* culture conditions. Additionally, when conidia from the same strain are plated as a lawn on solid medium hyphae grow closely together and are indistinguishable from each other (**Fig. 1B**). Interestingly, the *ΔarvA* strain colonies grow as dense micro-colony aggregates and do not form a colony biofilm (**Fig. 1B**). Dense aggregate formation is also observed at the air-liquid interface of submerged *ΔarvA* biofilms grown in liquid GMM (**Fig. 1B**).

**Figure 1.**
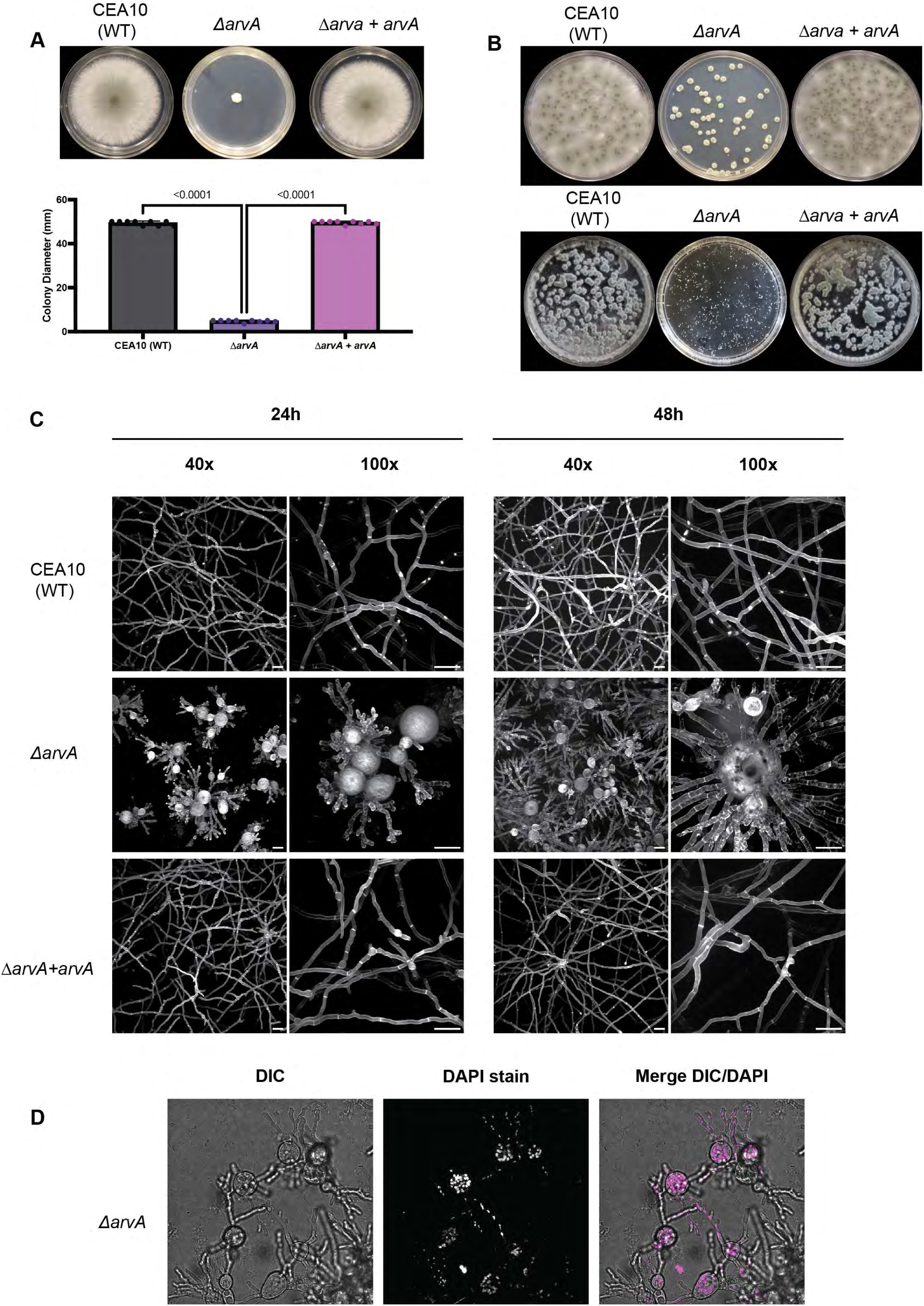
Loss of *Aspergillus fumigatus arvA* induces morphological changes on agar and in submerged colony biofilms. **A )** *ΔarvA* has significantly decreased hyphal growth in solid medium compared to wild-type (CEA10) and the reconstituted strain. 1×10^3^ conidia in 2 µL of CEA10, *ΔarvA* and *ΔarvA+arvA* strains were inoculated onto GMM plates and incubated for 72h at 37°C. 5% CO_2_. Colony diameter was measured using a ruler and reported as millimeters. Images and data are representative of 3 biological replicates with 3 technical replicates each. Statistical significance was determined using an ordinary one-way ANOVA with Tukey’s multiple comparison’s test. **B)** *ΔarvA* grows as micro-aggregates in solid and liquid media. The *ΔarvA* strain was inoculated onto solid (top panel, 50 ìL of a 1×10^3^ conidia/mL stock) or liquid (bottom panel, 1×10^5^ conidia/mL) GMM and incubated at 37°C, 5% CO_2_ for 72h for solid medium and 48h for liquid medium. The images are representative of multiple independent experiments **C)** *ΔarvA* has hyper-swollen conidia, loss of polarity, decreased hyphal growth and multiple germ tube emergence in submerged biofilms compared to wild-type and reconstituted strain. 1×10^5^ conidia/mL of CEA10, *ΔarvA* and *ΔarvA+arvA* were inoculated in filtered liquid GMM and incubated at 37°C, 5%CO_2_ for 24 and 48h. Biofilms were stained with 25 ìg/mL of calcofluor white (CFW) approximately 20 minutes prior to imaging. Images were taken on a Nikon spinning disk confocal microscope at 40X and 100X magnification, using a 405nm laser to visualize the CFW signal. Z-stacks were taken that encompassed the growth of the *arvA* null mutant, and max projections of those Z-stacks are shown. Scale bars are 50 µm. Raw images processed in FIJI as indicated in the methods. D) *ΔarvA* hyper-swollen conidia have multiple DAPI stained puncta. 1×10^5^ conidia/mL of *ΔarvA* was inoculated in filtered liquid GMM and incubated at 37°C, 5%CO_2_ for 24h. Biofilms were stained with 20 µg/µL of DAPI for 10 minutes prior to imaging. Images were taken on a Nikon spinning disk confocal microscope at 40X magnification, using a 405nm laser to visualize the DAPI signal. Z-stacks were taken that encompassed the growth of the *arvA* null mutant, and max projections of those Z-stacks are shown. Raw images processed in FIJI as indicated in the methods.

After observing the striking colony growth differences between *ΔarvA* and the parental and reconstituted strains, we assessed strain growth and morphology in a submerged biofilm model.

*A. fumigatus* dormant conidia size is approximately 2 µm and upon the start of germination conidia isotropically swell to ∼5 µm (28). After swelling, one germ tube develops from which hyphal growth begins. A second germ tube develops on the conidia on the opposite side of the first one, and continued hyphal growth develops a biofilm on a surface. At 24 hours, CEA10 and *ΔarvA+arvA* exhibit normal biofilm development at 40x magnification, and there are only two germ tubes per spore when assessing morphology at 100x magnification (**Fig. 1C**). Loss of *arvA (ΔarvA*) drastically alters submerged biofilm morphology. When compared to the parental and reconstituted strains, *ΔarvA* 24h biofilms have significantly decreased hyphal length (**Fig. 1C**).

Most striking, conidia of the *ΔarvA* strain are hyper swollen and a closer examination of the conidia at 100x magnification revealed multiple germ tubes emerging per conidia with hyphal hyperbranching indicating a loss of polarity (**Fig.1C**). At 48h, the Δ*arvA* strain continues to exhibit hyper-swollen conidia, a multitude of germ tubes per conidia, and hyphal hyperbranching that are not observed in CEA10 and *ΔarvA+arvA* (**Fig. 1C**). Combined, these results suggest that ArvA is necessary for the appropriate transition from isotropic to polarized growth in *A. fumigatus* and consequently wild-type biofilm formation.

These drastic changes in morphology observed in *A. fumigatus ΔarvA* have not been reported in yeast, therefore the existing fungal literature on Arv1 failed to provide a clear explanation for *ΔarvA* hyper-swollen conidia phenotype in submerged biofilms. However, human Arv1 has been observed to play a role in cell cycle telophase progression and the lack of Arv1 in human cells results in multinucleate cells (8). An accumulation of nuclei as a result of cell cycle progression perturbation could explain the size increase observed in *ΔarvA* conidia. We therefore hypothesized that the large size of the hyper-swollen *ΔarvA* conidia is in part due to an expansion in the number of nuclei. To test this hypothesis, we stained 24 hour Δ*arvA* submerged biofilms with DAPI. In support of the hypothesis, *ΔarvA* hyper-swollen conidia contained multiple distinct DAPI stained puncta suggestive of increased nuclei within each hyper-swollen conidia (**Fig.1D**). From these data, we conclude that *A. fumigatus* ArvA is critical for normal cell cycle progression and conidia germination under the conditions examined.

### ΔarvA displays differential susceptibility to cell wall and cell membrane stressors

After establishing that *ΔarvA* in *A. fumigatus* is an important determinant of growth and orphology, we wanted to test if reported phenotypes of *Δarv1* strains in other fungal species were conserved in *A. fumigatus*. One pathogenicity related phenotype attributed to the loss of Arv1 is a loss of thermotolerance (11, 29, 30). To test whether *A. fumigatus ΔarvA* exhibited an inability to grow at elevated temperatures, we employed an agar-based colony biofilm assay (**Fig. S1**). We inoculated 1×10^3^ conidia of CEA10, *ΔarvA* and *ΔarvA+arvA* onto GMM plates and incubated for 72h at 25°C, 37°C or 45°C with 5% CO_2_. CEA10 and *ΔarvA+arvA* strains were able to grow at each respective temperature with no growth or morphological differences (**Fig. S1**). In contrast, *ΔarvA* is only able to grow at 25°C and 37°C but not at 45°C (**Fig. S1**).

However, *ΔarvA* growth at 25°C did not look markedly different compared to 37°C. This result confirms a role for *A. fumigatus* ArvA growth at temperatures well above those found in mammals, and that growth and morphology aberrations exist in the absence of ArvA at both ambient and mammalian core body temperatures.

As growth at high temperatures requires alterations to membrane sterol and lipid composition, morphological and growth aberrations associated with loss of ArvA could be due to its predicted role as a lipid transporter. As ArvA has previously been shown to play a role in sterol and sphingolipid metabolism, we hypothesized that loss of *arvA* would alter susceptibility to membrane-targeting antifungals. Due to the striking morphological differences in modes of growth exhibited by Δ*arvA* relative to its parental (CEA10) and reconstituted strain (*ΔarvA+arvA),* we employed two different assays to assess susceptibility to cell wall and cell membrane stressors. Furthermore, as the cell membrane directly impacts cell wall homeostasis, we hypothesized that loss of *arvA* would also result in altered susceptibility to cell wall perturbing agents. We first used an agar-based colony biofilm assay to assess differential susceptibility to amphotericin B (cell membrane stressor), voriconazole (cell membrane stressor), calcofluor white (cell wall stressor) and caspofungin (cell wall stressor). We inoculated 1×10^3^ conidia of CEA10, *ΔarvA* and *ΔarvA+arvA* onto GMM plates treated with either amphotericin B (0.75 µg/mL), voriconazole (0.0625 µg/mL), caspofungin (0.125 µg/mL), or calcofluor white (25 µg/mL) and incubated for 72h at 37°C, 5% CO_2_ (**Fig. 2A**). Radial growth of each plate was measured and normalized to percent of their own untreated control. Treatment with amphotericin B reduced growth by ∼25% in both CEA10 and *ΔarvA+arvA* strains, and by ∼19% in *ΔarvA* with no statistical difference between the 3 strains observed (**Fig. 2A-B**). In contrast, a sub-MIC concentration of voriconazole (0.0625 µg/mL = 1/4 - 1/8 MIC in wild-type parental strain CEA10) reduced growth about 30% in CEA10 and *ΔarvA+arvA,* compared to ∼81% reduction in growth exhibited by *ΔarvA* (**Fig. 2A-B**). This increase in susceptibility to azoles in the absence of ArvA orthologs has been reported in *Saccharomyces cerevisiae* and *Candida albicans* (6, 12, 26).

**Figure 2.**
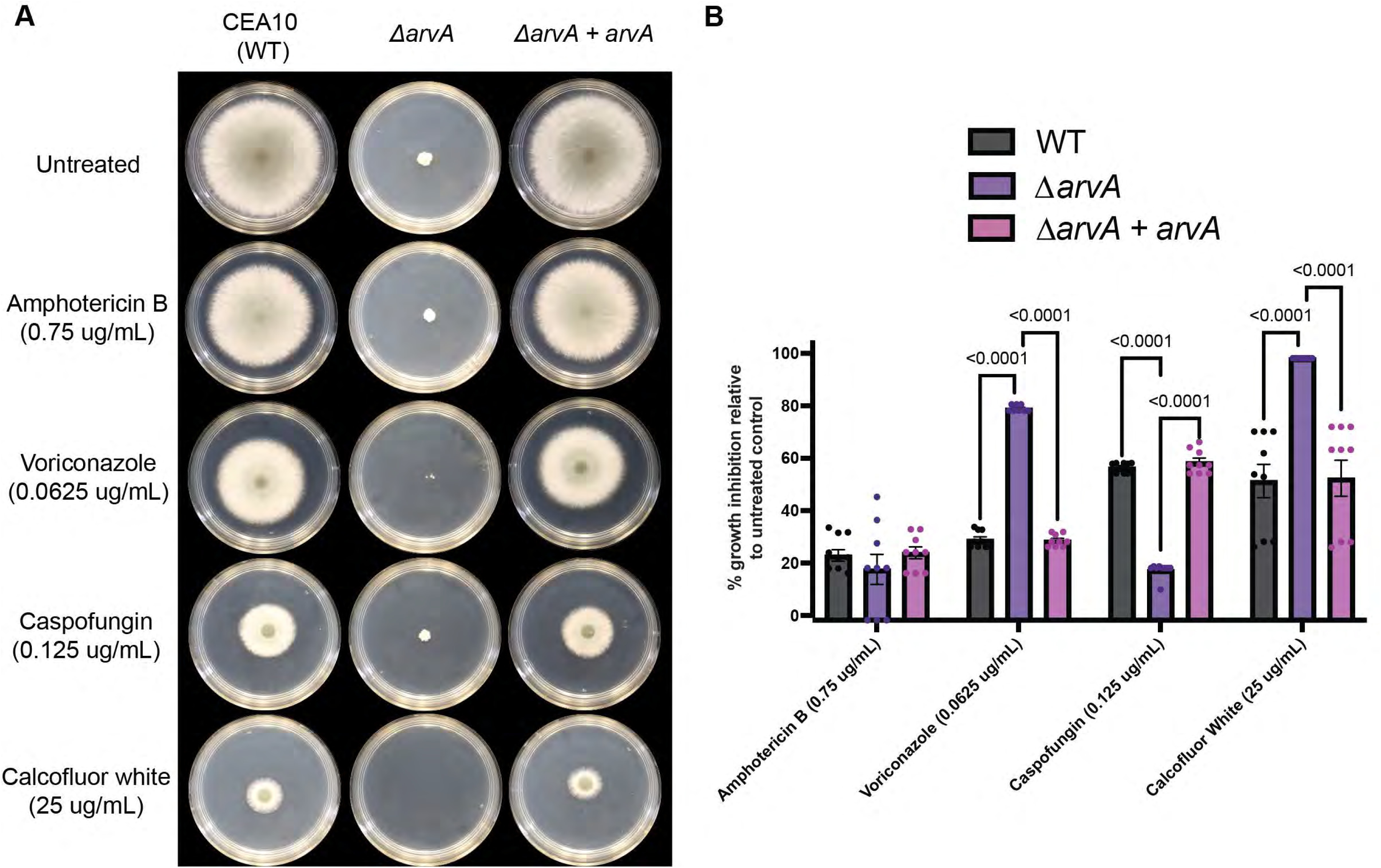
Loss of *A. fumigatus arvA* alters susceptibility to cell membrane and cell wall targeting antifungal agents. **A-B )** *ΔarvA* is more susceptible to voriconazole and calcofluor white and less to caspofungin relative to wild-type and reconstituted strains in agar-based colony assays. 1×10^3^ conidia in 2 µL of CEA10, *ΔarvA* and *ΔarvA+arvA* strains were inoculated onto GMM plates treated with amphotericin B (0.75 µg/mL), voriconazole (0.0625 µg/mL), Caspofungin (0.125 µg/mL), or calcofluor white (25 µg/mL) and incubated for 72h at 37°C. 5% CO_2_. Colony diameter was measured using a ruler and normalized to % growth inhibition relative to its untreated control. Images and data are representative of 3 biological replicates with 3 technical replicates each. Statistical significance was determined using an ordinary one-way ANOVA with Tukey’s multiple comparison’s test.

We next tested whether *ΔarvA* had altered susceptibility to the cell wall stress agents calcofluor white (CFW), a chitin binding molecule, and caspofungin, a β-glucan synthase inhibitor. Caspofungin inhibited CEA10 and *ΔarvA+arvA* growth ∼58% and ∼60% respectively, compared to only ∼19% growth inhibition of *ΔarvA* (**Fig. 2A-B**). The *ΔarvA* strain was therefore less susceptible to caspofungin mediated β-glucan synthesis inhibition. This result suggests that *ΔarvA* has changes in cell wall composition and/or the stress response to echinocandin treatment. In support of the *ΔarvA* strain having a different cell wall composition in comparison to parental and reconstituted strain, *ΔarvA* was hyper-susceptible to the chitin binding agent calcofluor white (CFW) (**Fig. 2A-B**). CFW inhibited CEA10 and *ΔarvA+arvA* strains growth ∼53-54% respectively compared to a striking 100% growth inhibition of *ΔarvA* (**Fig. 2A-B**). Overall, Δ*arvA* has altered susceptibility to voriconazole, caspofungin and CFW compared to its parental and reconstituted strain. However, these assays are potentially limited by the small radial growth exhibited by the *ΔarvA* strain (∼5mm) on solid agar, therefore any small change in radial growth when treated with a drug is amplified when normalized to the untreated strain.

Consequently, to determine if these finding are generalizable to another fungal growth model, we tested the impact of these stress agents using the *A. fumigatus* submerged biofilm model and an XTT based cell damage assay. As XTT readouts may be proportional to total fungal biomass, we first determined that a 34h *ΔarvA* biofilm had equivalent biomass to 16h CEA10 and *ΔarvA+arvA* biofilms and utilized these time points in subsequent assays (**Fig. 3A**). We grew Δ*arvA* (34 hours), CEA10 (16 hours) and *ΔarvA+arvA* (16 hours) biofilms in liquid GMM prior to treatment with either amphotericin B (0.5, 1 µg/mL), voriconazole (1 µg/mL), caspofungin (0.125, 0.25 µg/mL) or CFW (12.5, 25 µg/mL) for 3 hours. To assess the effect of drug treatment, metabolic activity (XTT reduction) was normalized to percent reduction in metabolic activity relative to untreated control (biofilm damage) (2). Treatment with 0.5 µg/mL amphotericin B led to a ∼62% and ∼68% reduction of metabolic activity in CEA10 and *ΔarvA+arvA* biofilms respectively compared to a ∼52% reduction in *ΔarvA* (**Fig. 3B**). This modest decrease in metabolic activity reduction observed in the *ΔarvA* strain was aggravated and became statistically significant with an increased concentration of amphotericin B (1 µg/mL) (**Fig. 3B**). Treatment with 1 µg/mL of amphotericin B reduced metabolic activity by ∼90 and ∼92% in CEA10 and *ΔarvA+arvA* strains respectively, while it reduced metabolic activity ∼72% in the *ΔarvA* strain (**Fig.3B**). These data suggest loss of *arvA* confers a modest decrease in amphotericin B susceptibility in a submerged biofilm model. With regard to the triazoles, voriconazole was not effective against biofilms of the parental strain CEA10, as previously observed (2, 31). In this assay, voriconazole only reduced the metabolic activity ∼7 and 12% of CEA10 and the reconstituted strain *ΔarvA+arvA* respectively (**Fig. 3C**). Voriconazole also did not reduce metabolic activity of the *ΔarvA* strain in this biofilm model (**Fig. 3C**). This is in contrast to what we observed in our agar-based colony biofilm assays (**Fig. 2C**). Taken together, these data suggest potential perturbations in the cell wall and/or lipid composition of the *ΔarvA* that impact antifungal drug susceptibility under specific conditions.

**Figure 3.**
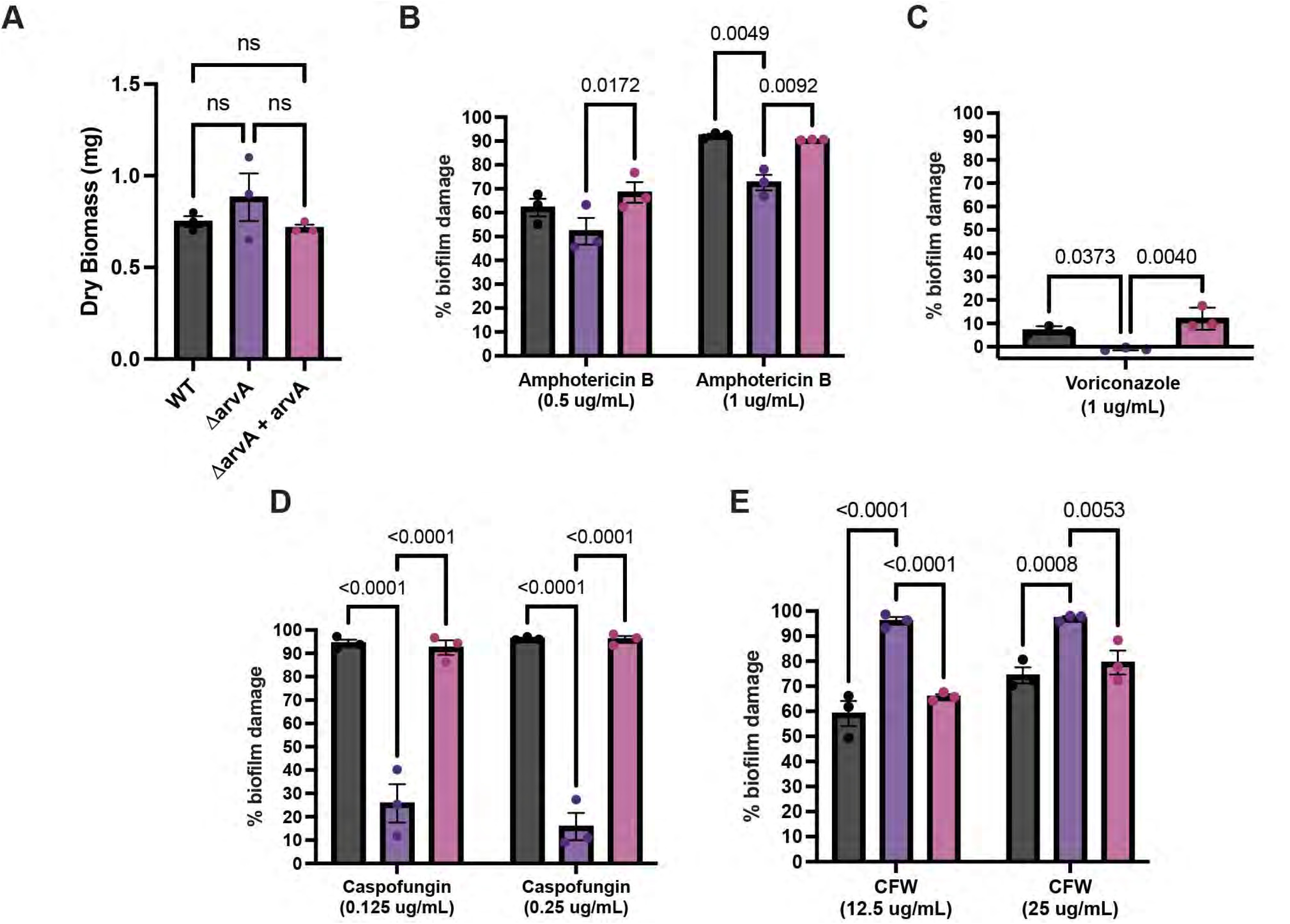
Loss of *A. fumigatus arvA* alters susceptibility to cell membrane and cell wall targeting antifungal agents in submerged biofilms. **A)** *ΔarvA* 34h biofilm biomass is equivalent to a 16h CEA10 and *ΔarvA+arvA* biofilm. 1×10^5^ conidia/mL of each strain was inoculated onto 6-well plates and incubated for 34h (*ΔarvA*) or 16h (CEA10, *ΔarvA+arvA*) at 37°C. 5%CO_2_. Biofilm biomass was determined by collecting biomass from 6-well plates into a tube, washed two times with water, lyophilized and weighed. Data are representative of 3 bio replicates, with 2-3 technical replications each. One-way ANOVA with multiple comparisons. **B-E)** *ΔarvA* is more susceptible to voriconazole and calcofluor white and less to Caspofungin relative to wild-type and reconstituted strains in submerged biofilms as measured by an XTT assay. 1×10^5^ conidia /mL of CEA10, *ΔarvA* and *ΔarvA+arvA* strains were inoculated into liquid GMM and incubated for 34h (*ΔarvA*) or 16h (CEA10, *ΔarvA+arvA*) at 37°C. 5%CO_2_. Media was removed and biofilms were treated with (voriconazole (0.0625 µg/mL), amphotericin B (0.75 µg/mL), calcofluor white (25 µg/mL), caspofungin (0.125 µg/mL) or vehicle control in liquid GMM for 3h. After treatment media was removed and metabolic activity was determined using XTT as described in the methods. To assess the effect of drug treatment, metabolic activity was normalized to percent reduction in metabolic activity relative to untreated control (biofilm damage). Data is representative of the mean of each technical replicate from three biological replicates. Statistical significance determined using 2-way ANOVA with Tukey’s multiple comparison’s test.

With regard to the echinocandins, treatment with the β-glucan synthesis inhibitor caspofungin was highly efficacious against the parental and reconstituted strains at the time points analyzed. 0.125 µg/mL of caspofungin reduced metabolic activity by 93% and 94% in CEA10 and *ΔarvA+arvA,* compared to only ∼26% reduction in the *ΔarvA* strain. Treatment with a higher dose of caspofungin showed a similar trend with reducing ∼96% of metabolic activity in CEA10 and *ΔarvA+arvA*, compared to only ∼16% in *ΔarvA.* This result confirms what we observed in agar-based colony biofilm plate assays (**Fig. 3D**) and supports that Δ*arvA* is less susceptible to β-glucan synthesis inhibition by caspofungin. Loss of *arvA* also increases susceptibility to calcofluor white (CFW). A low dose of CFW reduced ∼60% and ∼66% metabolic activity in CEA10 and *ΔarvA+arvA* (**Fig. 3E**). Strikingly, the same dose of CFW reduced ∼96% of metabolic activity in *ΔarvA* biofilms (**Fig. 3E**). This increased susceptibility to CFW in *ΔarvA* was replicated with 25 µg/mL of CFW treatment (∼97% reduction) (**Fig. 3E**). The *arvA* dependent susceptibility to CFW replicated what we observed in agar-based colony biofilm assays (**Fig. 2A-B**), further supporting that loss of ArvA impacts the composition and/or function of the fungal cell wall.

### Loss of arvA induces changes in cell wall polysaccharide exposure

To test whether Δ*arvA* has changes in cell wall polysaccharide exposure, we imaged and measured chitin and β-glucan cell surface exposure by staining with wheat germ agglutinin (WGA) and soluble Dectin-1 respectively (32, 33). We grew biofilms (1×10^5^ conidia/mL) in liquid GMM for 24h prior to staining with FITC-WGA (5µL/mL) and fixing with 4% paraformaldehyde. Biofilms stained with soluble Dectin-1-Fc (sDectin-1) were fixed prior to staining as indicated in the methods. Each strain with either WGA or sDectin-1 staining was imaged and images were processed and normalized using FIJI. Due to the morphological differences of *ΔarvA* (hyper- swollen conidia), we measured chitin and β-glucans of hyphae for appropriate comparison with the parental and reconstituted strain.

Both WGA and sDectin-1 were able to stain all three strains (CEA10*, ΔarvA, ΔarvA + arvA*) (**Fig. 4**). Interestingly, the Δ*arvA* strain exhibited both WGA and sDectin-1 signals (fluorescent aggregates) in the medium, suggesting potential cell wall shedding (**Fig. 4**). This phenotype may suggest that the *ΔarvA* cell wall is more fragile than the WT and reconstituted strain and is more susceptible to mechanical disruption in the staining and fixing process. As expected, CEA10 and *ΔarvA+arvA* showed no statistical difference in either WGA or sDectin-1 (**Fig. 4**).

**Figure 4.**
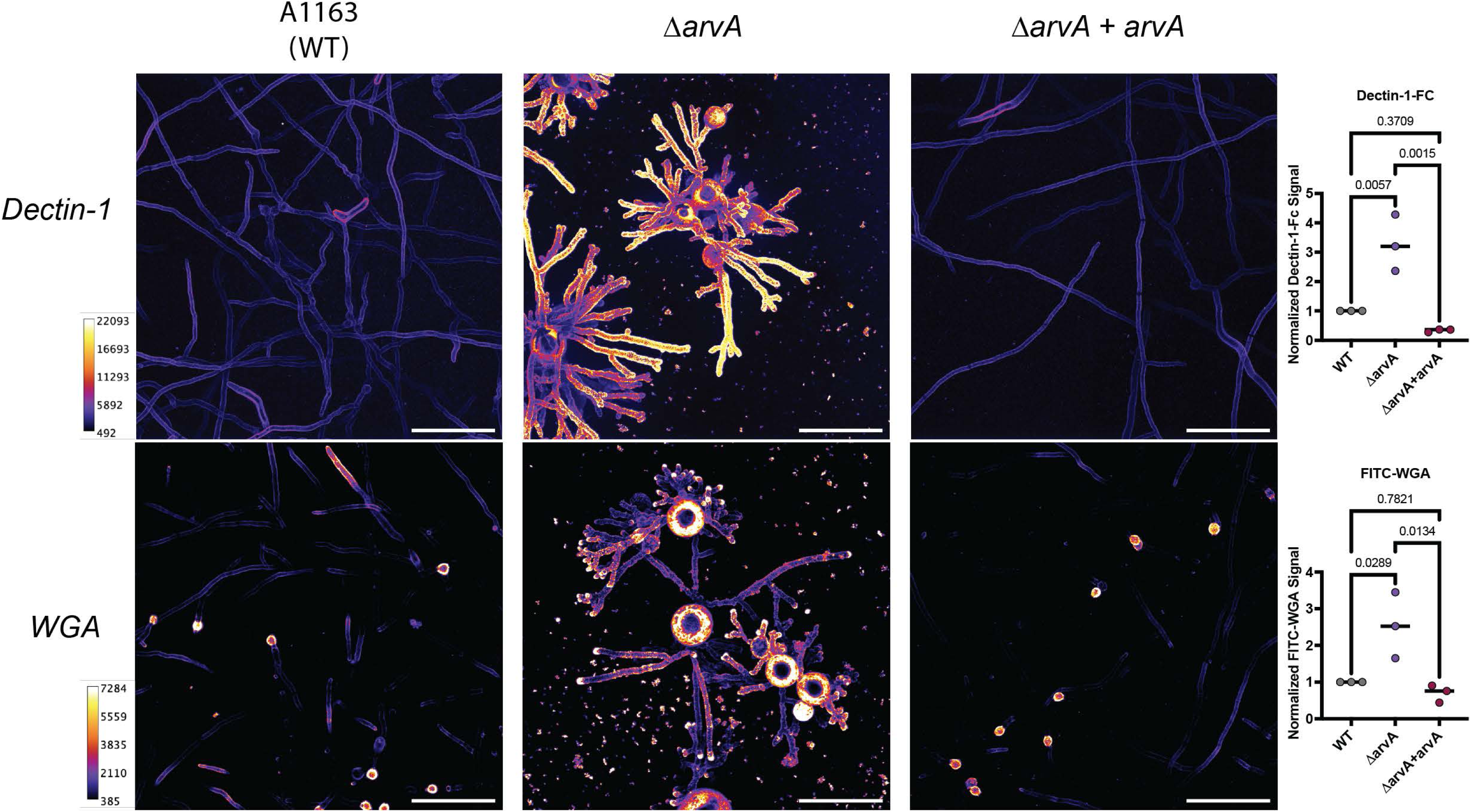
Loss of *arvA* shows increased exposure of chitin and β-glucans. 1×10^5^ conidia/mL of CEA10, *ΔarvA* and *ΔarvA+arvA* strains were inoculated onto filtered liquid GMM in 8-well Ibidi plates and incubated for 24h at 37°C, 5% CO_._ For WGA staining, biofilms were stained with 5µL/mL of FITC-WGA, incubated at room temp for 30 min, and subsequently washed with PBS. After washing, biofilms were fixed with 4% paraformaldehyde for 15 min. Soluble Dectin-1 staining was achieved by fixing biofilms with 4% paraformaldehyde for 15 min and stained with 5µg/mL of sDectin-1-Fc before incubating with Alexa Fluor 488 anti-human IgG (ThermoFisher). Biofilms were imaged and levels of exposed chitin and β-glucan measured as described in the methods. Quantification data is representative of 3 biological replicates. Each biological replicate (n=3) consists of 4 measurements on 3 different fields of view, for a total of 12 independent measurements of mean grey value per biological replicate. Scale bars are 50 µm. Statistical significance determined using an ordinary one-way ANOVA with Tukey’s multiple comparison’s test. Heatmap indicates intensity values of images.

WGA staining in CEA10 and *ΔarvA+arvA* strains was more prominent in the hyphal tips and in the conidia, while the Δ*arvA* strain showed a more equivalent staining throughout the entire fungal mass (**Fig. 4**). Additionally, WGA staining in *ΔarvA* is heavily punctate throughout the hyphae rather than the smooth, well-defined pattern observed on CEA10 and *ΔarvA+arvA* strains or even in sDectin-1 staining in *ΔarvA* (**Fig. 4**). This result suggests that *ΔarvA* has altered chitin localization in the hyphal cell wall. Overall, the Δ*arvA* strain had higher staining of both sDectin-1 and WGA compared to WT and the reconstituted strain (**Fig. 4**). Together these results suggest that the cell wall organization in *ΔarvA* hyphae differs from the wild-type and reconstituted strains.

### ΔarvA strain is fully virulent and can grow in vivo in mouse models of IPA

Given the significant *in vitro* growth and morphological defects of the Δ*arvA* strain we hypothesized that loss of ArvA would inhibit *A. fumigatus* pathogenesis and/or significantly reduce virulence in murine models of invasive pulmonary aspergillosis (IPA). To test whether *arvA* is necessary for pathogenicity and virulence, we challenged Triamcinolone (steroid) immune suppressed mice with the *arvA* isogenic strain set as previously reported (34, 35). On day 3 post fungal challenge (dpi 3), we sacrificed mice to evaluate histology (GMS, H&E) and quantify lung fungal burden. In separate experiments, pathogenicity and virulence were quantified utilizing a Kaplan-Meier survival curve analysis (**Fig. 5**).

**Figure 5.**
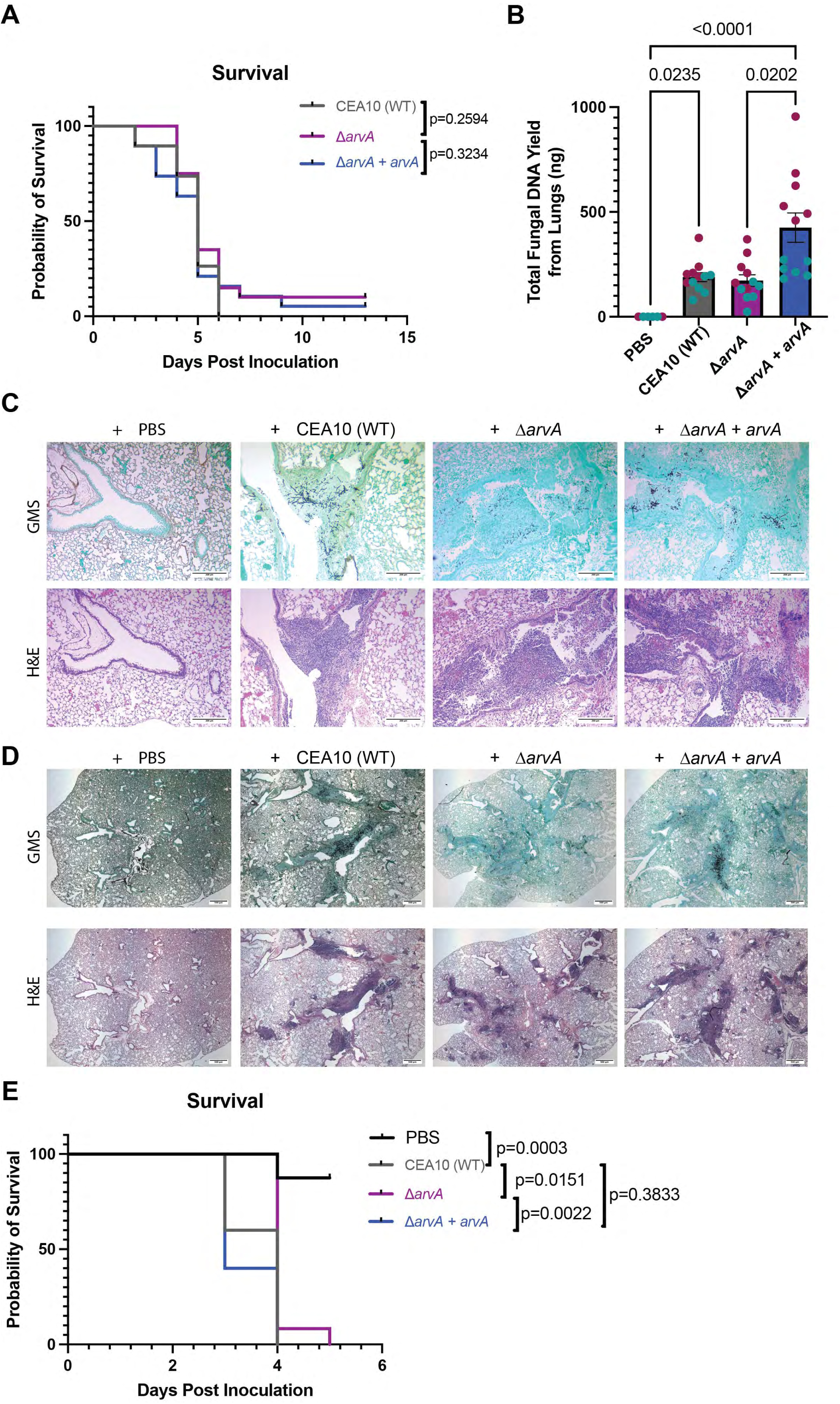
*ΔarvA* is fully virulent and able to grow in the lungs but shows reduced tissue invasion in mouse models of IPA. **A )** *ΔarvA* has no significant difference in murine survival relative to wild-type and reconstituted strains in immunosuppressed steroid mouse model of invasive pulmonary aspergillosis. Outbred female CD-1 mice 22-24g (Charles River Laboratories) were injected subcutaneously with 40mg/kg Kenalog-10 (triamcinolone acetonide; Bristol-Myer Squibb, Princeton, NJ). 24hr after steroid injection, mice were inoculated intranasally with 1×10^5^ conidia (WT-CEA10, Δ*arvA* or Δ*arvA* + *arvA*) in 40µL sterile PBS while under isoflurane anesthesia. Data are represented on a Kaplan-Meier plot to determine significance (18-20 mice/group from two independent experiments). **B)** *ΔarvA* fungal burden is similar to wild-type in murine lungs. Mice were injected with Kenalog-10 and inoculated as described above. At 3 days post inoculation, select mice were euthanized and lungs were excised from the chest cavity, and genomic DNA extracted for 18S rDNA qPCR quantification of fungal DNA. A one-way ANOVA with a Krukal Wallis test comparing all groups was used to determine significance (12 mice per group from two independent experiments). **C)** *ΔarvA* shows reduced tissue invasion compared to wild-type and reconstituted strains. Mice were injected with Kenalog-10 and inoculated as described above. At 3 days post inoculation, 4 mice per strain were euthanized, cannulated, and lungs were inflated with 10% buffered formalin phosphate. Lungs were paraffin-embedded and stained for hematoxylin and eosin (H&E) and Gomori methenamine silver (GMS). H&E and GMS stained lungs were imaged at 10x objective using a standard upright light microscope fitted with an AmScope MU1000 camera. Scale bars were added using AmScope calibration slide and ImageJ software (ImageJ2, v. 2.14.0/1.54f). Scale bars are 200μm. Data are representative of 6-8 mice per group from two independent experiments. **D)** Experimental details as in Figure 5C except images taken at 2.5X and scale bar=500μm. **E)** *ΔarvA* is fully virulent in a leukopenic murine model of IPA. 6-7 week old female CD-1 mice were injected with 150mg/kg cyclophosphamide (IP) day -2 and 40 mg/kg Kenalog-10 subcutaneously day -1. 24hr later they were inoculated intranasally with 1×10^5^ conidia/40µL PBS of the indicated strains. Survival was monitored over the course of 5 days. A Kaplan-Meier Survival analysis and Log-Rank test comparing WT CEA10 to the other three groups was used to determine significance between isogenic sets. For PBS, n=8; for WT and recon, n=10, and for *ΔarvA*, n=12.

Surprisingly, the *ΔarvA* strain showed no significant difference in survival in this IPA murine model compared to CEA10 and *ΔarvA+arvA* strains (**Fig. 5A**). Furthermore, *ΔarvA* and CEA10 had no significant difference in fungal burden as measured by qPCR from mouse lungs at 3 dpi (**Fig. 5B**). Given the likely increase in nuclei observed *in vitro,* we wondered if the qPCR assay might predict higher *arvA* null mutant strain fungal burden than is actually present in the lung.

Histopathology examination suggests that the *ΔarvA* strain grows in the mouse lung though it appears to be less robust than the parental and reconstituted strains (**Fig. 5C**). However, in spite of its stunted morphology *in vitro*, the *ΔarvA* strain exhibits no quantifiable evidence of a morphological difference *in vivo* compared to CEA10 and the reconstituted strain *ΔarvA+arvA* (**Fig. 5C**). Interestingly, the histopathology of the Δ*arvA* strain also suggests the strain is largely contained within the larger airways and shows an apparent reduction in invasion into surrounding parenchyma, with some evidence of accumulation in the aveolar spaces (**Fig. 5C**).

In contrast, CEA10 and *ΔarvA+arvA* strains are able to invade the lung parenchyma and proliferate (**Fig. 5C**). Further assessment of the histology slides at 2.5X magnification confirmed that the *ΔarvA* strain is largely restricted to the larger bronchioles with a modest presence in the alveolar spaces (**Fig. 5D**).

Consequently, histopathology suggests mortality mechanisms may be different in this IPA model with *ΔarvA* compared to the parental and recon strain. One possibility is that the *ΔarvA* strain induces hyperinflammation due to the changes in cell wall morphology, and potential shedding of cell wall material, observed *in vitro*. Increased exposure of *A. fumigatus* β1,3-glucans is correlated with higher inflammation as a result of increased immune response *in vivo* (36, 37). To test whether the mortality observed in *ΔarvA* challenged steroid treated mice is due in part to immunopathogenesis, we conducted a pathogenesis experiment using a chemotherapeutic induced leukopenic murine model of IPA (**Fig. 5E**). Interestingly, similar to the steroid murine model, the *arvA* strain was fully virulent, although mortality was delayed by one day compared to WT and *ΔarvA*+arvA infected mice. The mortality delay was statistically significant, however, as all *ΔarvA*-challenged mice succumbed to infection, its relevance for therapeutic development is likely low. We conclude that immunopathogenesis differences at least partially contribute to *ΔarvA*-induced pathogenicity in the steroid model, though other factors are likely in play to fully explain host mortality. Taken together, these data surprisingly suggest that loss of *arvA* does not substantially impact disease progression and mortality in two immunologically distinct murine models of IPA, and beg the question why the severe *in vitro* growth phenotype is not recapitulated *in vivo*.

### In vitro *A. fumigatus* growth requirements in absence of ArvA

We subsequently hypothesized that the ill-defined murine lung environmental conditions at least partially rescued the *ΔarvA in vitro* stunted growth and morphology. To try and identify what lung nutrient/condition might rescue *ΔarvA* morphology *in vivo*, we tested an array of different media (GMM, SCN, RPMI, and lung homogenate) and supplements (Ergosterol, Calcium, Myo-inositol). None of these conditions were able to rescue the Δ*arvA* growth and morphology we observed *in vitro* in GMM (**Fig. 6**). As we observed Δ*arvA* in some alveolar spaces in histopathology, we next tested growth between CEA10, *ΔarvA,* and *ΔarvA+arvA* in a natural calf pulmonary surfactant (used clinically) containing medium. As expected, culturing in surfactant reduced overall growth of CEA10 and *ΔarvA+arvA* strains compared to their growth in the relatively nutrient rich L-GMM, while the *ΔarvA* strain had modestly improved germination compared to L-GMM (**Fig. 6**). Furthermore, *ΔarvA* morphology in pulmonary surfactant looked more similar to CEA10 and *ΔarvA+arvaA* than in L-GMM (**Fig. 6**). While these data do not rule out other unknown host nutrients/conditions that promote pathogenicity and virulence in the absence of ArvA, they suggest that growth in a host relevant nutrient source, surfactant, is a better indicator of *ΔarvA* murine model outcomes.

**Figure 6.**
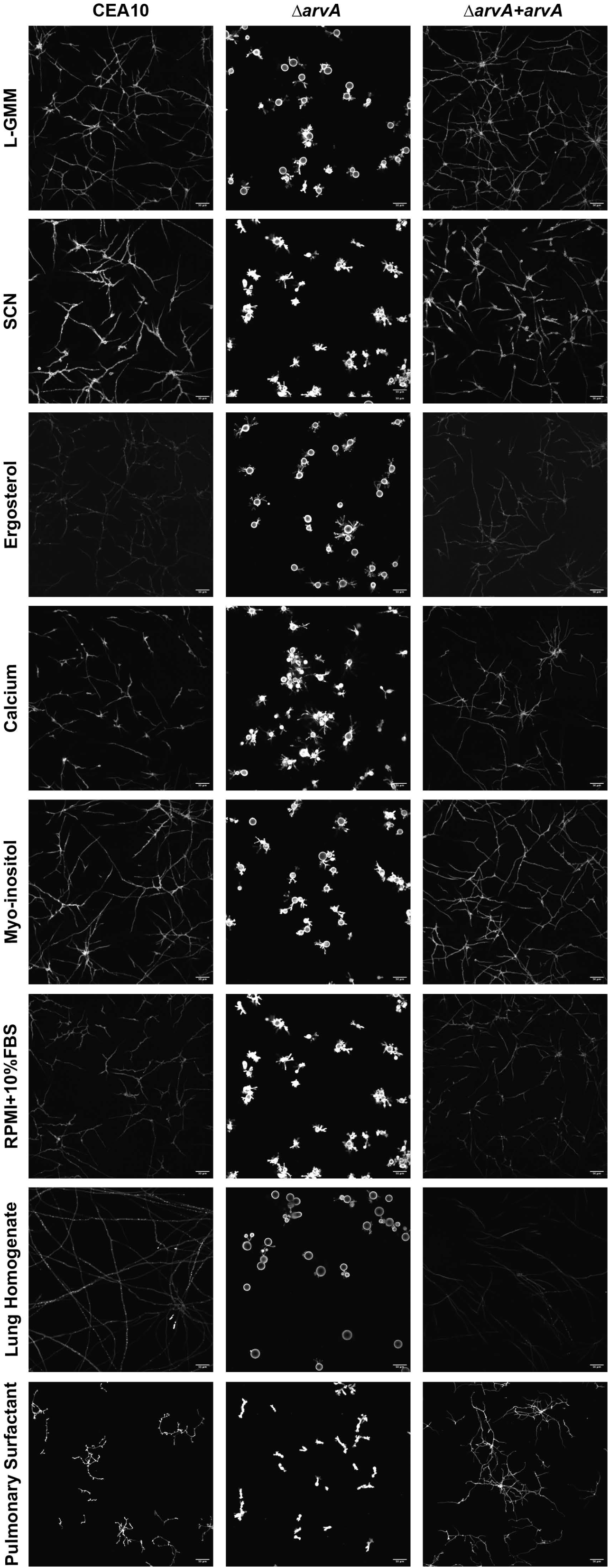
*ΔarvA* is partially rescued in natural calf pulmonary surfactant. CEA10, Δ*arvA* or Δ*arvA* + *arvA* were seeded into L-GMM, SCN, RPMI+10% FBS, lung homogenate, lung surfactant or natural calf pulmonary surfactant medias or L-GMM supplemented with calcium, myo-inositol, or ergosterol 1×10^5^/ml for 18hr. Biofilms were stained with Calcofluor white 10 µg/mL and imaged on a confocal microscope at 60X, scale bars=50µm. Data are representative of three biological experiments with 2 technical replicates each.

## DISCUSSION

ArvA is conserved across eukaryotes and is important for proper lipid distribution in many organisms though its precise molecular function remains unclear. In *A. fumigatus*, single gene replacement of the *arvA* gene resulted in stunted hyphal growth and visible changes in biofilm morphology in solid agar and liquid media assays. A hallmark of this stunted biofilm morphology is hyper-swollen, multinucleate conidia that continue swelling in spite of hyphal growth occurring (**Fig.1**). It is unclear why the *ΔarvA* mutant, conidia hyperswell, characteristic of isotropic growth, but simultaneously initiate multiple polarity axes as evidenced by the multile germ tubes that arise from each conidium. This is highly atypical for *A. fumigatus*, as conidia usually swell from ∼2 µM to ∼5 µM and stop swelling after polarity is established and the first germ tube emerges (38, 39).

The mechanism and reason behind the altered germination progression is unclear, but could be related to ArvA’s predicted function as an important lipid transporter associated with the endoplasmic reticulum. Membrane lipid organization and composition is important for the establishment and maintenance of polarized growth in other fungi including *Aspergillus nidulans* and the dimorphic fungal pathogen *Candida albicans* (40). In *A. nidulans* ceramide synthase mutants result in altered polarized growth morphologies and loss of the lipid transporter encoding gene *dnfA* results in altered hyphal and colony morphology (41, 42). In *C. albicans*, steep gradients of phospholipids facilitate polarized growth in both the budding yeast and filamentous form (43, 44). Additionally, lipid transporters, specifically Dsr2, are important for invasive growth and virulence (45). Taken together, these studies support the hypothesis that *A. fumigatus* ArvA is a potential lipid transporter involved in organizing lipids important for key fungal life cycle morphological transitions.

Along these lines, multiple DAPI stained puncta in the *A. fumigatus arvA* null mutant suggest potential cell cycle progression dysfunction in the absence of *arvA.* Multinucleate cells, as a result of loss of function of *arvA,* have not been reported in other fungi (5, 6, 12, 15). This phenotype, however, has been reported in human cells with reduced levels of the homolog ARV1 (8). In these cells, ARV1 plays a role in cell cycle progression by recruiting IQGAP1 to the cleavage furrow (8). IQGAP1 is a plasma membrane-associated protein that regulates the actin cytoskeleton and is needed for cell polarity (46). In the fungal pathogen *C. albicans*, downregulating its IQGAP1 homolog, IQG1, results in multiple polarity sites that cause multi-budding (47). As the *A. fumigatus ΔarvA* strain has hyper-swollen conidia with multiple DAPI stained puncta, and multiple germ tube emerging, we hypothesize that ArvA plays a role in cell cycle progression through interaction with IQGAP1 homologs in *A. fumigatus* similar to ARV1 in humans. This is an exciting and novel phenotype in fungi that could expand knowledge of cell cycle progression in *A. fumigatus*. Interestingly, ARV1 has been shown to translocate from the cleavage furrow to the intercellular bridge during the telophase stage of the cell cycle in human cells (8). Studying cellular localization of ArvA in *A. fumigatus* and potential interactions with the IQGAP1 homolog (Rho-family GTPase, AFUB_020720) is worthy of future investigations.

Further support of the importance of ArvA in *A. fumigatus* biology is evident by the altered antifungal drug susceptibility of the *arvA* null mutant. While the *ΔarvA* strain had a non- significant decrease in susceptibility to amphotericin B in the agar-based colony biofilm assays (**Fig. 2A-B**), in the XTT biofilm assay, Δ*arvA* had a surprising significant decrease in amphotericin B susceptibility (**Fig. 3B**). This result is surprising due to previous reports in *S. cerevisiae*, where a mutant *arv1* strain without the conserved AHD domain is hyper-susceptible to amphotericin B (12). However, the assays and mutants in our study and the *S. cerevisiae* study are different, potentially explaining the difference in susceptibility observed in the two studies. Regardless, the data indicate loss of ArvA likely alters ergosterol membrane content and thus assessing ergosterol in the cell membrane of *ΔarvA* biofilms in future studies is warranted.

With regard to triazoles, in the agar-based colony biofilm assay, *ΔarvA* in *A. fumigatus* has increased susceptibility to voriconazole (**Fig. 2A-B**), supporting previous reports of triazole susceptibility in *Δarv1* strain in *C. albicans* and *S. cerevisiae* (*6, 12, 26*). Azole susceptibility in *Δarv1* strains has been proposed to be due to Arv1p’s physical interaction with the azole target Erg11p, which stabilizes Erg11p and increases its half-life. Based on the azole susceptibility we observed with *A. fumigatus ΔarvA* , we hypothesize ArvA likely plays a similar role in *A. fumigatus.* Additionally, we showed that loss of *arvA* likely results in altered cell wall composition and PAMP exposure. Relative to the parental strain (CEA10) and reconstituted strain *(ΔarvA+arvA*), the Δ*arvA* strain has increased WGA staining suggesting an increase in chitin exposure. Furthermore, *ΔarvA* is hyper-susceptible to calcofluor white, which binds chitin in the cell wall. In other fungal species, Arv1 null mutants hyperaccumulate chitin and are hypersusceptible to calcofluor white (3, 14). As we did not measure chitin levels directly, we cannot conclusively say there is chitin hyper-accumulation in *ΔarvA* cell walls in *A. fumigatus*.

However, based on the increased WGA signal (**Fig. 4**) and the conserved hyper-susceptibility to calcofluor white (**Figs. 2 and 3**), which match observations in other fungal species, it is reasonable to speculate that there is increased chitin deposition in *ΔarvA.* Additionally, our microscopy showed irregular and punctate WGA staining in *ΔarvA*, suggesting that even though there is increased chitin in the cell wall, it is disorganized which may lead to the increased calcofluor white susceptibility. It is unclear how alterations in the *ΔarvA* impact antifungal drug susceptibility, and it is worth considering whether these alterations impact drug penetration into the cell.

In yeast, *Δarv1* has defects in β-glucan masking and increased levels of exposed β-glucans, even though the strain itself has reduced levels of both β1,6- and β1,3-glucans (48). In *A. fumigatus* we observed increased binding of dectin-1 to *ΔarvA* cells compared to the wild-type (**Fig. 4**), consistent with a higher level of exposed β-glucans in the absence of *arvA*. Furthermore, *ΔarvA* had decreased susceptibility to caspofungin, which, based on the findings in yeast, we postulate is due to impaired synthesis of β-glucans. Alternatively, the likely increased chitin observed in the absence of Arv may compensate for the loss of β-glucans. Interestingly, *ΔarvA* had cell wall fragments in the medium that stained with both WGA and dectin-1, suggesting cell wall instability and shedding. Altogether these results validate some of the published cell wall phenotypes of strains with loss of ArvA homologs in other fungi and introduce new questions regarding ArvA’s role in cell wall composition, assembly, and regulation of β-glucan synthesis in a filamentous fungus. As β-glucan synthase is localized to the cell membrane, it is possible that alterations in the membrane lipid content due to loss of ArvA lead to changes in β-glucan and chitin synthase activity and/or localization.

With significant growth and cell wall defects, it was expected that loss of ArvA would dramatically attenuate *A. fumigatus* pathogenicity and virulence. Consequently, perhaps the most surprising result of this study is the persistent pathogenicity and virulence of the *ΔarvA* strain in two immunologically distinct murine models of invasive pulmonary aspergillosis (IPA) (**Fig. 5**). The only observable differences in *in vivo* growth between *ΔarvA*, CEA10 and *ΔarvA+arvA* was a decrease in tissue invasion and perhaps modest increase in *ΔarvA* presence in the alveolar spaces. Though limited by the resolution of histology imaging, to the extent that we could observe with this approach, the severe *in vitro* morphological and polarity defects of *ΔarvA* were absent *in vivo*. Consequently, we were curious if the nutrient environment of the airways rescued *ΔarvA* morphology and growth. However, differences in growth and morphology between the wild-type and *ΔarvA* strains persisted across numerous different *in vivo* relevant nutrient sources that we examined in vitro (**Fig. 6**). Intriguingly, the one nutrient condition where we observed relatively similar growth and morphology among CEA10, *ΔarvA,* and *ΔarvA+arvA* was in natural calf lung surfactant. Importantly, surfactant did not fully rescue *ΔarvA* growth compared to CEA10 and *ΔarvA+arvA* growth in L-GMM, but rather significantly restricted growth of CEA10 and *ΔarvA+arvA* to better match *ΔarvA.* As this growth medium, which contains 35mg/ml phospholipids (including phosphatidylcholine) and surfactant proteins B and C (0.7mg/ml), is reflective particularly of the alveolar spaces in the lung, *in vivo* lung surfactant may be worth considering in the assessment of *A. fumigatus* mutants and strains as part of future studies. From this perspective, it is possible *ΔarvA* is more resistant to surfactant mediated fungal growth inhibition. For example, the surfactant protein D is known to restrict growth of *A. fumigatus* (49). It would be of further interest to test if a co-culture experiment with primary differentiated epithelial cells would also rescue the *ΔarvA* morphology as seen *in vivo*.

Taken together these data raise the question whether ArvA is a viable antifungal drug target for *A. fumigatus* due to the apparent *in vivo* complementation of *ΔarvA’s* severe growth defects.

Future studies should examine infection outcomes with *ΔarvA* during antifungal drug treatment to determine if increased efficacy of triazoles observed *in vitro* would occur in the *in vivo* infection microenvironment. Consequently, a significant take home message of this study is that the still ill-defined pulmonary nutrient environment of an *A. fumigatus* infection makes predictions of pathogenicity and virulence, and thus viability of a candidate gene product as a drug target, based on *in vitro* phenotypes highly uncertain.

## METHODS

### Generation of arvA mutant and reconstituted gene

The *arvA* mutant was generated using a dual Cas9-gRNA system as previously described for *A. fumigatus* (50) (all primers used listed in Table S1). Briefly, two crRNA spanning the *arvA* gene were designed (g158 - gacgaccgctctttgggcaa; g159 - ggtcgaatgcgtcgtcccgc), which were separately annealed to tracrRNA to generate gRNAs. The gRNAs were conjugated with Cas9 to generate the Cas9 RNP complex. The repair template was generated from a Hygromycin (Hyg) plasmid by PCR amplification using RAC 7051 and RAC 7052. Two micrograms of repair template and the Cas9 RNP complex were transformed into *A. fumigatus* protoplasts as previously described (51).

To reconstitute the *arvA* null mutant, a plasmid was generated by amplifying the *arvA* gene locus using the primers RAC 7317 and RAC 7318. PCR using RAC 7319 and RAC 7320 primers and a ptrA-Ampilcillin vector was then utilized to generate a product with overhangs complementary to RAC 7318 and RAC 7317, respectively. Using HiFi assembly, a plasmid was generated from the two PCR products such that the *ptrA* selection marker is adjacent to the 3’ UTR of *arvA*. For transformation, the repair construct was amplified using primers RAC 7321 and RAC 6321, which carry flanks with 35-40 bp homology adjacent to a gRNA (g65-TCTCCTTCATAAGCGACCAG) cut site at the *Aft4* transposon (52). Transformation into *arvA* mutant was performed as described above.

### Southern blot for verification of reconstituted strain copy number

A Southern blot was conducted as described previously (53). Briefly, 25 ìg of WT, mutant, and the reconstituted strain gDNA were digested overnight using the *Hind*III enzyme and separated on a 1% agarose gel. The bands were transferred onto a positively charged nylon membrane and were hybridized overnight with a DIG-labeled probe specific to *ptrA*. Post-hybridization, washing, and DIG detection were carried out according to manufacturer instructions (Roche).

### Agar-Based Colony Biofilm Drug Susceptibility Assays

1×10^3^ conidia in 2 ìL of WT-CEA10, *ΔarvA* (CEA10), and *ΔarvA + arvA* (CEA10) strains were inoculated onto glucose minimal media plates (GMM, 6 g/liter NaNO3, 0.52 g/liter KCl, 0.52 g/liter MgSO4·7H2O, 1.52 g/liter KH2PO4 monobasic, 2.2 mg/liter ZnSO4·7H2O, 1.1 mg/liter H3BO3, 0.5 mg/liter MnCl2·4H2O, 0.5 mg/liter FeSO4·7H2O, 0.16 mg/liter CoCl2·5H2O, 0.16 mg/liter CuSO4·5H2O, 0.11 mg/liter (NH4)6Mo7O24·4H2O, 5 mg/liter Na4EDTA, 1% glucose; pH 6.5, 1.5% agar) treated with Amphotericin B (0.75 µg/mL), voriconazole (0.0625 µg/mL), calcofluor white (25 µg/mL), or Caspofungin (0.125 µg/mL). Plates were incubated at 37°C, 5%CO_2_ for 72h unless otherwise indicated. Colony diameter was measured using a ruler and normalized to % untreated control. Each experiment has 3 biological replicates with at least 3 technical replicates each.

### Biofilm Drug Assays

CEA10, *ΔarvA*, and *ΔarvA + arvA* strains were grown for 16hr in liquid-glucose minimal media (L-GMM) in either 96 well plates (XTT assay) or 6 well plates (biofilm biomass) at 1×10^5^ conidia/ml L-GMM in 5% CO_2_ at 37°C. *ΔarvA* conidia were grown in L-GMM for 34 hours under the same conditions as CEA10 and *ΔarvA + arvA* strains to match biomass. In 96 well plate cultures, media was removed and fresh L-GMM with either vehicle control, voriconazole (Sigma), Caspofungin (Sigma), calcofluor white (VWR), or amphotericin B (Caymen Chemicals), was added for 3hr. Media was removed and metabolic activity was assessed by addition of 0.5 µg/ml XTT (2,3-Bis-(2-Methoxy-4-Nitro-5-Sulfophenyl)-2H-Tetrazolium-5-Carboxanilide; Thermo Scientific) plus 25 µM menadione (Enzo) for an additional hour, or until untreated control wells turned dark orange. Supernatants were transferred to a new plate and optical density (OD) was measured at 450nm. To assess the effect of drug treatment, metabolic activity was normalized to percent reduction in metabolic activity relative to untreated control (biofilm damage) (2). Biofilm biomass was determined by collecting biomass from 6-well plates into a tube, washed two times with water, lyophilized and weighed.

### Dectin-1 and WGA staining

Dectin-1 and WGA staining were done according to (54). Briefly, 200 ìL of 1×10^5^ conidia/mL in optically clear LGMM were added to an 8-well Ibidi slide at 37 °C for 24hrs. For WGA staining, biofilms were stained with 5 µL/mL of FITC-WGA in optically clean LGMM, incubated at room temp for 30 min, and subsequently washed with PBS. After washing, biofilms were fixed with 4% paraformaldehyde for 15 minutes. For biofilms stained with both WGA and CFW, CFW was added to fixed biofilms after WGA staining at a final concentration of 25 µg/mL.

Dectin-1 staining was achieved by fixing biofilms with 4% paraformaldehyde for 15 minutes. After fixation, biofilms were washed with PBS and then blocked for 20 minutes using RPMI supplemented with 10% fetal calf serum and 0.025% Tween 20. Blocking solution was removed and 5 µg/mL of Dectin-1-Fc in blocking solution was added to the biofilms and was incubated at room temperature for 1hr. Biofilms were washed with PBS and a 1/300 dilution in PBS of Alexa Fluor 488 anti-human IgG (ThermoFisher) was added and incubated at room temperature for 1hr. After this step, biofilms were washed with PBS and maintained in PBS until use.

### Microscopy

All samples were imaged on a Nikon spinning disk confocal microscope equipped with a Yokogawa CSU-W1 spinning head using a 40, 60 or 100x oil objective as indicated. To visualize FITC-WGA and Dectin-1 FC/Alexa Fluor 488, a 488nm laser was used. To visualize CFW and DAPI a 405nm laser was used. Images were processed in Nikon Elements and Fiji software (55, 56). Images were acquired using z-stacks to capture the volume of the sample and the same exposure time and laser intensity were used for all images within an experiment.

### Analysis of WGA and Dectin-1 fluorescent signal

To account for the different morphologies exhibited by *ΔarvA* in spore size, we quantified WGA and Dectin-1 on the hyphae. Images were analyzed using Fiji (55, 56). Using a line with a thickness of 10 in Fiji we measured the intensity of the stained cell wall at single z-slice which is positioned in the center of the cell. We repeated this process in multiple hyphae per image. We then found the minimum (background) and maximum (peak) intensities and subtracted the average background intensities from the individual peak intensities per image.

### Microscopy to assess changes in ΔarvA morphology with different media

To perform screening for conditions that rescue cell morphology of the *arvA* mutant strain, 1×10^4^ conidia/mL were grown in the media indicated in either a standard 96 well plate for panel A, or 1×10^5^ conidia/mL inoculated into Ibidi glass-bottom slide dishes (B,C, and D) (Cat.No: 80827) for 24 hours at 37°C, 5% CO_2_. For panels B, C, and D, cells were stained with 25 ìg/mL of calcofluor white (CFW) approximately 30 minutes prior to imaging. Images were taken on a Nikon spinning disk confocal microscope using a 20x objective, using a 405nm laser to visualize the CFW signal, except for in the ammonium condition, where DIC static images were taken due to interference with CFW signal in. Z-stacks were taken that encompassed the growth of the *arvA* mutant, and max projections of those Z-stacks are shown (raw images processed in FIJI). Images in (A) were taken using a camera attached to a standard benchtop light microscope using a 40x objective.

### Media preparation for ΔarvA morphology experiments

Glucose minimal media (GMM) (As described in *Agar-Based Colony Biofilm Drug Susceptibility Assays)* was supplemented with 500 ìg/mL myo-inositol or CaCl2 25mM. Myo-inositol was resuspended in water, filter sterilized, and added to autoclaved GMM. For ergosterol supplementation, 2.5mg ergosterol first resuspended in a 1mL solution of 1:1, ethanol: Tween80, and added 1:100 to sterile GMM. SCN was generated by adding 10 g/L glucose, 6 g/L NaNO_3_, 0.502 g/L KCl, 0.50 2g/L MgSO_4_ 7H_2_O, 1.52 g/L KH_2_PO_4_, 0.025 g/L MnCl_2_ 4H_2_O, 2.2 mg/L ZnSO_4_ 7H_2_O, 1.1 mg/L H_3_BO_3_, 0.5 mg/L MnCl_2_ 4H_2_O, 0.16 mg/L CoCl_2_ 5H_2_O, 0.16 mg/L CuSO_4_ 5H_2_O, 0.5 mg/L FeSO_4_ 7H_2_O, 5 mg/L Na_4_EDTA and pH adjusted to 6.5 and autoclaved. After autoclaving, the following sterile solutions were added: L-Glutamine solution, to a final concentration of 146.14 mg/L, SC mix (Sunrise Science Catalog#1300-030) to 1g/L, NaNO_3_, to 101.864 mg/L). RPMI1640 with L-Glutamine was purchased from Corning (product number 10-040-CV). RPMI + low glucose: HyClone RPMI + 2.05mM L-Glutamine Cat No: SH30011.04, 10.4 g/LD-Glucose, 2 g/L. Fetal bovine serum was added to RPMI 1640 to a final concentration of 10% (FBS+RPMI). To generate lung homogenate medium, murine lungs were isolated, homogenized, and resuspended in 4mL PBS. Natural calf surfactant (Infasurf®; final concentration 100ìg/ml) was made by diluting in water with trace elements (2.2g/L, ZnSO4·7H2O,1.1g/liter H3BO3, 0.5 mg/liter MnCl2·4H2O, 0.5 mg/liter FeSO4·7H2O, 0.16 mg/liter CoCl2·5H2O, 0.16 mg/liter CuSO4·5H2O, 0.11 mg/liter (NH4)6Mo7O24·4H2O, 5 mg/liter Na4EDTA).

### Microscopy to assess changes in ΔarvA morphology with pulmonary surfactant

Strains were seeded with 1×10^5^ conidia/mL for each indicated strain. Strains were grown in the indicated media in 24-well Ibidi plates for 24 hours at 37°C, 5% CO_2_. After growth, Biofilms were stained with Calcofluor White 10ug/ml PBS and fluorescent images were taken on a Nikon Spinning Disc Confocal Microscope. Images were taken using a 405nm laser.

### Murine Model Studies

Mice were randomly allocated into autoclaved cages at 3-4 mice per cage with HEPA-filtered air with food and water available ad libitum. For the steroid model, outbred female CD-1 mice 22-24g (Charles River Laboratories) were injected subcutaneously with 40mg/kg Kenalog-10 (triamcinolone acetonide; Bristol-Myer Squibb, Princeton, NJ). 24 hours after Kenalog injection, mice were inoculated intranasally with 1×10^5^ conidia (CEA10, *ΔarvA* or *ΔarvA + arvA*) in 40 ìl sterile PBS while under isoflurane anesthesia. For the leukopenic model, mice were injected with cyclophosphamide 150mg/kg intraperitoneally 48 hours prior to and 72 hours post fungal inoculation. Mice were treated with Kenalog, inoculated, and monitored as described in the steroid model. For both models, mice were monitored three times per day for 14 days for signs of disease. Mortality data were analyzed with a Kaplan-Meier Survival Curve to determine significance between each group. All studies were performed under strict accordance with the Guide for the care and use of Laboratory Animals (57). All protocols used were approved by the Institutional Animal Care and Use Committee (IACUC) at Dartmouth College.

### Histopathology

CD-1 mice were housed, treated, and inoculated as described above. At 3 days post inoculation, n=4 mice/strain were euthanized, canulated, and lungs were inflated with 10% neutral buffered formalin. Lungs were stored in 10% neutral buffered formalin for a minimum of 24 hours, then sectioned and stored in 70% ethanol until paraffin embedding. Paraffin-embedded lung sections were stained for hematoxylin and eosin (H&E) and Gomori methenamine silver (GMS). H&E and GMS stained lungs were imaged at 40x objective using a standard upright light microscope fitted with an AmScope MU1000 camera. Scale bars were added using AmScope calibration slide and ImageJ software (ImageJ2, v. 2.14.0/1.54f).

### Fungal Burden

CD-1 mice were housed, treated, and inoculated as described above. At 3 days post inoculation, select mice were euthanized and lungs were excised from the chest cavity, rinsed in sterile PBS, and flash frozen in liquid nitrogen. Lungs were then lyophilized and bead beaten with 2.3mm Zirconia beads. Genomic DNA was extracted according to the manufacturer’s protocol (E.Z.N.A. fungal DNA mini kit; Omega Biotech) and modifications were made on the protocol as previously described (54). qPCR quantification of fungal DNA was performed as previously described (58).

### Ethics Statement

Animal studies were strictly carried out in accordance with the recommendations in the Guide for the Care and Use of Laboratory Animals (59). The animal experimental protocol 00002167 was approved by the Institutional Animal Care and Use Committee (IACUC) at Dartmouth.

## DATA AVAILABILITY

All underlying data are available in either the supplement or through an author’s request.

## ACKNOWLEDGEMENTS

This work was funded by NIAID/NIH R21/R33AI140878 (RAC), NIAID/NIH R01AI181215, and NIAID/NIH R01AI130128. CGP was supported by the John H. Copenhaver, Jr. and William H. Thomas, MD 1952 Fellowship at Dartmouth. Core facility support provided by NIH grant P20-GM113132 to the Dartmouth BioMT COBRE. Additional support was provided by the Cystic Fibrosis Foundation Research Development Program (STANTO19R0) and NIH P30-DK117469 (Dartmouth Cystic Fibrosis Research Center). AJ and NEK were supported by an NIH/NHBLI T32HL134598 grant (O’Toole, George). NEK is also supported by a Ruth L. Kirschstein NIH/NIAID F31AI188661 award. SV is supported by a post-doctoral fellowship award from the Cystic Fibrosis Research Foundation. The authors would also like to thank Dr. Daniel Aridgides, Dartmouth Health, for providing the natural calf pulmonary surfactant, Ann Lavanway in the Dartmouth College Microscopy core for assistance with imaging, and the CCMR staff and veterinarians at Dartmouth for animal husbandry.

